# A role for reward valuation in the serotonergic modulation of impulsivity

**DOI:** 10.1101/2020.01.17.910778

**Authors:** Stephanie S. Desrochers, Emma Lesko, Valerie M. Magalong, Peter D. Balsam, Katherine M. Nautiyal

## Abstract

Impulsive behavior is a deleterious component of a number of mental health disorders but has few targeted pharmacotherapies. One contributing factor to the difficulty in understanding the neural substrates of disordered impulsivity is the diverse presentations of impulsive behavior. Defining the behavioral and cognitive processes which contribute to different subtypes of impulsivity is integral to understanding and treating disorders with dysregulated impulsive behavior. Our approach was to first determine what behavioral and cognitive phenotypes are associated with increased impulsive behavior, and then probe if they could causally contribute to increasing impulsivity. We used a mouse model for disordered impulsivity - mice lacking the serotonin 1B receptor (5-HT_1B_R) which have deficits specific to impulsive action, and not other components of impulsive behavior. Here we report, that in addition to increased impulsive action, mice lacking expression of 5-HT_1B_R also have increased goal-directed responding and motivation, with no differences in extinction, development of habitual behavior, delay discounting, or effort-based discounting. Interestingly, mice lacking 5-HT_1B_R expression did show an overall increase in the choice of higher value rewards, increased hedonic responses to sweet rewards, and responded more to cues that predict reward, compared to controls. We developed a novel paradigm to demonstrate that increasing anticipated reward value could directly increase impulsive action. Furthermore, we found that 5-HT_1B_R KO-induced impulsivity could be ameliorated by decreasing the reward value relative to controls, suggesting that the increased 5-HT_1B_R-associated impulsive action is a result of increased reward valuation. Taken together, these data show that the effects of serotonin on impulsive action are mediated through the modulation of hedonic value, which may alter the reward representations that motivate action. Additionally this work supports a role for reward valuation as an important substrate in impulsive action which may drive clinically-relevant increases in impulsivity.

## Introduction

Impulsivity is an important component of daily life, but can lead to many deleterious outcomes such as making unhealthy eating decisions or excessive online shopping. It is also a core component of a number of psychiatric disorders including attention deficit disorder, alcohol and substance use disorders, and gambling disorder [1–3]. Treatment options to decrease impulsivity are limited, and those that exist are not targeted to impulsive behavior. One underlying issue in the development of approaches to reduce impulsive behavior lies within the complexity of the broad construct of impulsivity. Individual facets of what is broadly referred to as impulsive behavior, for example impulsive action (e.g., acting on a whim) and impulsive choice (e.g., wanting immediate gratification), are likely mediated by different behavioral/cognitive processes with different neurobiological substrates [3–6]. In fact, some argue against the use of the term impulsivity at all given the divergence and independence of its latent factors, in favor of more specific labels which have internally consistent behavioral and neurobiological substrates [7]. Consistent with this view, processes such as reward valuation, compulsivity, motivation, attention deficits, novelty-seeking, and anxiety have all been associated with some aspects of impulsivity using trait-level behavioral measures in humans (particularly in psychiatric populations) and in preclinical models [8–14]. Understanding how these behavioral and cognitive substrates are causally associated with different components of impulsive behavior will lead to an understanding of the behavioral/cognitive scaffolding and associated underlying neural circuits which lead to dysregulated impulsivity.

Ours and others’ previous work has examined how different dimensions of impulsivity can be dissociated, behaviorally and biologically [4,15,16]. Specifically, impulsive action, characterized by the reduced ability to withhold or delay responses, is independent from impulsive choice, which includes an exaggerated discounting of future or risky rewards. While many studies have focused on the role of dopamine in the modulation of impulsivity, serotonin signaling is particularly relevant when focusing on dissociation of the neurobiology of impulsive choice from impulsive action. Manipulation of serotonin signaling in humans and rodents supports its role in modulating impulsive action specifically [5,17–19]. The mechanisms of these effects are likely through a number of the 14 serotonin receptors including 5-HT_1B_, 5-HT_2A_, and 5-HT_2C_. We previously reported that mice lacking the 5-HT_1B_ receptors show increased impulsive action, but not impulsive choice [4,20–22]. Interestingly, in humans, 5-HT_1B_R has also been implicated in disorders which present with dysregulated impulsivity, such as substance use and gambling disorders. Cocaine-dependent individuals had reduced 5-HT_1B_R activation compared to controls [23], and single-nucleotide polymorphisms in *htr1b* were associated with cocaine, alcohol, and heroin abuse [24]. In preclinical models, genetic knockout of 5-HT_1B_R caused increases in cocaine self-administration in mice [25], and administration of a 5-HT_1B_R agonist decreased behavior motivated by both cocaine and sucrose rewards [26].

Serotonin signaling could act through several cognitive and behavioral mechanisms which may promote difficulty withholding responses (elevated impulsive action). While it is common to attribute this type of impulsivity to a deficit in inhibitory control, behavioral action is a product of both inhibitory and excitatory processes. Given intact inhibitory control mechanisms, an exaggerated representation of reward value may also make “holding back” difficult due to increased drive, resulting in increased impulsivity. The representation of reward value that motivates behavioral responding arises from experiences with the current hedonic value of rewards [27–29]. When response-outcome contingencies are learned, the likelihood of responding is guided by the incentive value of the outcome, which can be seen in the hedonic reaction [30]. For example, when subjects are hungry, food rewards have greater value than when subjects are sated, and thus hedonic reactions are diminished. When the expected reward value is lowered, motivation is also reduced. While the hedonic reactions to reward referred to as “liking” and the motivational processes that energize behavior referred to as “wanting “ can be dissociated, wanting is generally directed at available outcomes that are liked [31–34]. Here we refer to the increased responding for an appetitive outcome as reward reactivity to include components of both liking and wanting.

Both increased reward reactivity and increased impulsivity are found in patients with substance use disorders, and potentially represent risk factors for the development of addictive disorders [35–39]. In this framework, substance use disorders may arise from deficits in inhibitory control, but also from exaggerated hedonic valuation and/or through increased incentive salience, as in the Incentive-Sensitization Theory [40]. It is possible that the contribution of the 5-HT_1B_R to drug seeking behaviors is due to the modulation of hedonic reactions to drugs, at least in part. Reward reactivity has also been implicated in gambling disorder, with increased pleasure derived from winning potentially promoting the escalation of the behavioral addiction [41,42]. Additionally, in gambling disorder patients, levels of 5-HT_1B_R in brain regions associated with reward processing, including the ventral striatum, correlated with the severity of the disorder [43]. These considerations suggest the possibility that 5-HT_1B_R may impact impulsivity through changes in hedonic reactions and thus alter the representation of the value of a reward that motivates behavioral action.

The goal in the present studies was first to investigate what behavioral/cognitive processes, such as reward valuation, contribute to deficits in behavioral inhibition, and second to understand which process mediates the effect of serotonin on impulsive action. Specifically, we explored the effect of 5-HT_1B_R on potential substrates of impulsivity including goal-directed responding, motivation, habitual-like responding, and hedonic responding. While each of these can be conceptualized as unique behavioral phenotypes with distinct neural substrates, we focused on how alterations in impulsive action could potentially be subserved by changes in these processes. We specifically tested the hypothesis that the influence of serotonin on impulsivity is mediated by effects on the valuation of reward outcome. Additionally, by assessing these phenotypes in a mouse model for pathological impulsivity with deficits limited to impulsive action (absence of 5-HT_1B_R), we were also able to determine how associated behavioral mechanisms are related to different domains of impulsivity [5]. Coming to a better understanding of the specific neural *and* behavioral substrates of different dimensions of impulsivity will help us understand how these components combine to generate dysregulated impulsivity in psychiatric disorders.

## Methods

### Mice

Animals were bred in the Center for Comparative Medicine at Dartmouth College, or in the Department of Comparative Medicine at the New York State Psychiatric Institute. All mice were weaned at postnatal day (PN) 21 into cages of 2-5 same sex littermates on a 12:12 light-dark cycle, and maintained on ad lib chow until experimental operant behavioral testing began at 10-14 weeks. The floxed tetO1B mouse model was used to generate groups of mice lacking expression of 5-HT_1B_R through crosses to a βActin-tTS mouse line (tetO1B+/+ females crossed to tetO1B+/+::βActin-tTS+ males), as previously reported [20]. In the validation of the Variable Value Go/No-Go paradigm, only tetO1B+/+ control mice were used. In all other studies, tetO1B:: βActin-tTS+ mice and their littermate controls-tetO1B:: βActin-tTS-mice were used. For the adult rescue groups, tetO1B:: βActin-tTS+ mice were fed chow with doxycycline (DOX; 40mg/kg, BioServ) beginning at PN60 in order to rescue expression of the 5-HT_1B_R in the adult mouse, as previously validated and reported [20]. All procedures were approved by the Institutional Animal Care and Use Committees of the New York State Psychiatric Institute or Dartmouth College.

A summary table of the mice used in these experiments is provided in Table S1. One group of male (N=23) and female (N=35) mice were used in goal-directed behavior, extinction, concurrent choice, and satiety-induced devaluation. One mouse was excluded from progressive ratio due to technical issues, and one mouse died prior to the test, resulting in N=22 males and N=34 females included in the final analysis for the progressive ratio. Subsets of the total group were used in satiety-induced devaluation experiments (males N=23, females N=18) and Go/No-Go and delay discounting (males N=12, females N=8). One mouse was excluded from delay discounting due to not meeting criteria (see delay discounting methods below). A separate group of mice was used to test effort-based discounting (males N=7, females N=14). Two mice were excluded from effort-based discounting due to not meeting criteria. A naïve group of mice (N=19, all female) were used to test free consumption of evaporated milk diluted at different concentrations. Additional groups of naïve mice were used in the lickometer (males N=6, females N=5) and Pavlovian-to-instrumental transfer (PIT; males N=6, females N=9) studies. One mouse was excluded from the PIT study because of a failure to lever press. An additional naïve group of 12 control mice (males N=7, females N=5) was used for the validation study of the novel Variable Value Go/No-Go paradigm. Finally, for the study of the role of the 5-HT_1B_R in the Variable Value Go/No-Go paradigm, an additional group of naïve mice (males N=14, females N=7) was used to test the effect of 5-HT_1B_R expression. Three mice were removed due to not meeting criteria during lever training. A subset of the animals from this experiment (males N=13, females N=5) were used to examine the effect of 5-HT_1B_R expression on chow consumption.

### Operant Behavioral Apparatus

Operant studies were conducted in eight identical chambers (Med Associates, St. Albans, VT) individually enclosed in ventilated isolation boxes. Each operant chamber consisted of stainless steel modular walls, and stainless steel bar floors. Each chamber contained a noseport receptacle for the delivery of liquid reward by a dipper (0.02ml cup volume), with head entry detected by an infrared beam break detector. On either side of the noseport, the chamber contained two ultra-sensitive retractable stainless steel levers placed 2.2 cm above the chamber floor. In paradigms in which only one of the two levers was used, the lever was counterbalanced across mice and remained the same throughout all paradigms.

There were LEDs located above each lever, and a houselight and speaker located on the upper portion of the wall opposite the levers. A computer equipped with MED-PC IV (Med Associates Inc., St Albans, VT) computer software delivered stimuli and collected behavioral data.

### Operant Behavioral Training

Operant training and testing were run 5-7 days a week. Mice were maintained at approximately 90% of their free-feeding weight. Water was provided ad libitum throughout the experiment. Undiluted evaporated milk was used as the reward for all operant studies in MedAssociates chambers. All mice were first trained to retrieve an evaporated milk reward through head entry into the receptacle, and then trained to press one of the two retractable levers to receive the evaporated milk reward on a continuous reinforcement (CRF) schedule. Daily sessions ended when mice received a maximum of 60 rewards, or after 60 minutes elapsed if the maximum had not been reached. Mice were trained until the criterion of 55 lever presses in a 60 minute session was reached. The mice were then trained on a random ratio (RR) schedule of escalating effort requirements (3 days of RR-5, 3 days of RR-10, 3 days of RR-20). The data from the last day on each schedule was analyzed. Subsequently, they were tested in extinction trials, concurrent choice, satiety-induced devaluation (a subset), and then progressive ratio. A subset of mice were then tested in Go/No-Go and delay discounting paradigms.

### Progressive Ratios of Responding

Following random ratio testing, mice were tested on a progressive ratio (PR) schedule for three consecutive days. A PRx2 schedule was used in which the number of lever presses required to receive a reward doubled following each reward. The session ended following either 2 hours, or a 3 minute period in which no lever presses were recorded [44]. The total number of lever presses summed over the session. The total number of lever presses rather than break point was used in the factor analysis to provide a continuous rather than categorical variable. One mouse was excluded from analysis due to technical problems with the operant box.

### Extinction Testing

Mice were exposed to an RR-20 schedule of reinforcement for 3 days, before being tested in 3 consecutive days of extinction training. Mice were placed in the operant box with the lever extended, however rewards were not administered. Lever presses and head entries were recorded for the duration of the 60 minute extinction sessions.

### Concurrent Choice

Following 3 days of RR-20 schedule of reinforcement, mice were placed in the operant box on each of 2 days with either freely available chow pellets or freely available evaporated milk in a cell culture dish. The lever of the operant box was also extended and was rewarding the mice with evaporated milk on a RR-20 schedule. These chow and milk conditions were counterbalanced across mice over the 2 days separated by a no choice RR-20 schedule day. Chow pellets and the dish of evaporated milk were weighed before and after the test session. Lever presses and head entries were recorded during the 60 minute session.

### Satiety-Induced Devaluation

Following 3 days of RR-20 schedule of reinforcement, mice were prefed either chow or evaporated milk on each of 2 days, counterbalanced across mice. Mice were placed individually in a holding cage similar to their home cage for 1h, and were free to consume an unlimited amount of either chow or evaporated milk presented in a cell culture dish. Chow pellets, the dish of evaporated milk, and the mice were weighed before and after the hour-long prefeeding session. Mice were then placed in the operant box and allowed to lever press for a RR-20 schedule of reinforcement. Lever presses and head entries were recorded during the 60 minute session.

### Food Consumption

Mice were temporarily housed in individual cages for measurement of food consumption with ad lib access to water. Mice were placed on the food restriction protocol 48h prior to testing in the “food restricted” state testing, to mimic the food restriction state of the operant paradigms which consisted of 1.5h free access to chow daily. 24h following the 1.5h free access, chow was returned to mice and intake was measured at 1h, 3h, and 24h time points. Following this 24h ad lib period, mice continued to have free access to chow for an additional 48h prior to “sated” state testing, when intake was recorded for a 24h period.

### Go/No-Go

Mice were trained and tested as previously described [20]. Briefly, following training on Go Trials, mice were presented with 7 daily sessions consisting of 30 discrete Go trials and 30 No-Go trials which were pseudo-randomly presented across blocks of 10 trials with a variable ITI averaging 45s. In No-Go trials, the lever was presented simultaneously with 2 cues (the house lights turning off, and a small LED light above the lever turning on). A lever press during the 5 second trial caused the lever to retract, the house lights to turn on, the LED light to turn off and a new ITI to begin without any reward for that trial. A lack of presses during the 5 sec trial resulted in a reward presentation. The impulsivity index was calculated by subtracting the proportion of correct No-Go trials from the proportion of correct Go trials.

### Delay Discounting

Mice were trained and tested as previously described [4]. Briefly, following training mice were presented with two levers for which presses resulted in either small or large (3x volume) rewards. The large reward was assigned to the lever which was initially least preferred by the mice, and remained consistent throughout the paradigm. Each daily session began with 10 forced choice trials (five on each lever randomly distributed) to ensure a minimum experience with each lever in each session, before presentation of 20 experimental choice trials. Mice were trained in 14 sessions with no delays on either lever. One mouse was eliminated because it did not meet the criteria of greater than 25% preference for the large lever averaged over the last 3 sessions. Subsequently, a delay was introduced after the large reward lever was pressed, before the reward was presented. There was no delay for the small reward and time delays for the large reward (0, 2, 4, 6, 8, or 10 s) were presented in separate sessions with 3 days for each time delay, in ascending delay order. Data were used from the last session of each time delay. A linear equation was fit to the preference data for each mouse over all delays, and the slope, intercept, and indifference point (imputed delay at 50% preference) was calculated from the linear regression.

### Effort Discounting

Mice were initially trained as described for the delay discounting paradigm by presenting two levers for which presses resulted in either small or large (3x volume) rewards. The large reward was assigned to the lever which was initially least preferred by the mice, and remained consistent throughout the paradigm. Each daily session began with 10 forced choice trials (five on each lever randomly distributed) to ensure a minimum experience with each lever in each session, before presentation of 20 experimental choice trials. Mice were trained in 14 sessions, after which two mice were eliminated because they did not meet the criteria of greater than 25% preference for the large lever averaged over the last 3 sessions. Subsequently, the fixed ratio (FR) schedule was increased from FR1 for the large reward lever, with 3 days at each of the follow schedules: FR2, FR4, FR8, FR16, FR24, FR32. The small reward lever remained at the FR1 schedule throughout the paradigm. Any single press to the small reward lever resulted in presentation of the small reward and termination of the trial. Percent preference for the large reward was calculated as the percentage of choice trials in which the large reward was obtained. Data from the last session at each FR schedule are presented and used for statistical analysis.

### Consumption of Varied Value Rewards

Prior to testing in this paradigm, mice were previously exposed to evaporated milk in both consumption tests and 13 weeks of operant behavioral testing under food restriction (as described above) rewarded with 100% evaporated milk in a variety of reinforcement paradigms (data not shown). For the reward testing, mice were placed individually in a cage and given 5 minute free access to a small cell culture dish (Falcon, 35mmx10mm) with varying concentrations of evaporated milk in a separate clean cage identical to their home cage. Milk concentration was varied across 5 days of testing, with 33%, 66%, 100%, 66%, and 33% on each day respectively (data was only analyzed for first 3 days because of anchoring effects on the descending concentration presentation). Mice were weighed immediately before and after testing to determine milk consumption during the session. Because of the potential inaccuracies in weighing the dishes due to milk spillage or bedding being pushed into the dishes, change in mouse weight was used to assay consumption over this short 5 minute timeframe.

### Lickometer

A Davis Rig 16-bottle Lickometer (Med Associates MED-DAV-160M) was used to test the effect of 5-HT_1B_R expression on reward reactivity to various concentrations of sucrose in sated and restricted conditions as described previously [45]. Mice were water restricted for 5 days of initial training, during which mice were placed individually in the apparatus and allowed to drink water for 30 min from the spout which recorded licks using a capacitance-based system. Subsequently, mice were maintained on *ad libitum* water, except for the night before exposure to a new concentration of sucrose to promote maximal consumption for habituation to the new taste. Mice were exposed daily to sucrose in the testing chamber in a number of conditions, and licking behavior was recorded for 30 minutes. The order of exposure was: 10% sucrose with water restriction (1 day), 10% sucrose with food restriction (2 days), 10% sucrose sated (2 days). These conditions were then repeated in the same order with 2% sucrose. For food restricted conditions, mice were food deprived from the previous days’ testing, and given 1h free access to food following testing. For sated conditions, mice had ad lib access to food and water for at least 24h. Conditions were run for two consecutive days to measure stability of licking within each set of parameters. There were no differences between any two days within the same condition, so data was averaged across the two days for analysis. Number of licks over the whole session, and lick rate for the first 2 minutes were analyzed. The first two minute lick rate was used as a way to assess hedonic component of licking behavior without the influence of post-ingestion satiety-related factors [46,47].

### Pavlovian-to-instrumental transfer

Mice were tested in a modified Pavlovian-to-instrumental transfer protocol aimed at assessing the extent to which a Pavlovian conditioned stimulus (CS) can support the acquisition of a novel instrumental response, as previously described [48]. All mice were first trained to retrieve an evaporated milk reward through head entry into the reward receptacle of the Med Associates chambers for 5 sessions. Mice were then trained for 12 sessions in a Pavlovian conditioning phase in which a cue (conditioned stimulus, CS) was paired with an evaporated milk reward. In each session, mice experienced 20 CS+ presentations (10s tone or white noise) in which a dipper containing evaporated milk reward was presented 5s following the cue onset. In each session, mice also experienced 20 presentations of a CS− with which no reward was associated. CSs were presented in a pseudo-random order, with variable ITIs averaging 60s (30-90s range). The conditioned stimuli of either a tone or white noise were counterbalanced across mice. The number of nosepokes into the reward receptacle was analyzed during CS+ and CS− presentations for all sessions with the immediately preceding 10s of ITI responding subtracted out (elevation score). There was no instrumental conditioning phase, and so the instrumental transfer test was performed on the day following the 12^th^ Pavlovian conditioning session. In the transfer test session, mice were presented with two levers extended for 45 min. A drop of evaporated milk reward was placed on each lever to promote lever pressing. Lever presses resulted in a 3s presentation of either the CS+ or CS−, but no reward was presented. The CS paired with the left or right lever was counterbalanced across mice and CS type. The number of presses to each lever was recorded, and grouped by association with CS+ or CS− across mice. The difference score (CS+ minus CS− lever presses) was calculated for each mouse.

### Variable Value Go/No-Go Paradigm

To assess the effect of reward value on impulsive action, we developed a novel paradigm based on the Go/No-Go Test of impulsive action. Mice were trained as described in Operant Behavioral Training, except CRF training took place with both levers extended such that pressing either lever provided reward. All mice initially sampled each lever. For the validation study, training continued for 6 days, by which point all mice had formed stable and strongly biased lever preferences, which was determined based on average percentage of presses during the final three days of CRF training (range: 77% to 100%). For the experimental study, training continued for 7 days, with 3 mice being excluded from future testing due to not acquiring lever pressing behavior. The remaining mice again formed biased lever preferences (range: 61% to 100%). In order to cause a reversal of their preference, the less preferred lever was then rewarded with three times the amount of evaporated milk reward compared to the more preferred lever, which remained at 0.02ml evaporated milk. In order to deliver the larger, 0.06ml reward, the dipper was activated three times in short succession, as previously described. In these reversal sessions, mice were presented with 10 forced choice trials (5 per lever) in which only one lever was extended until the lever was pressed (requiring them to sample each lever), followed by 20 choice trials in which both levers were presented. Mice were required to reach a criterion of 25% choice for the higher reward lever (averaged over the final 3 days of training) in order to be included in future testing. After 14 sessions, mice in the validation study were choosing the high reward lever 69 ± 6 % of the time (averaged over the final 3 days of training). In the experimental study, 3 mice failed to reach the 25% criterion and were removed. With this exclusion, mice chose the higher reward lever 57 ± 3 % of the time (averaged over the final 3 days of training). 60 trials were presented in each session, with 30 trials presented on each of the large and small reward levers randomly in blocks of 10 trials. In all trials, the lever extended for 5 sec. A press within 5 seconds initiated reward delivery, and lever retraction (“Successful Go Trial”). Otherwise, the lever retracted after 5 seconds and no reward was delivered (“Unsuccessful Go Trial”). Finally, mice were exposed to 8 sessions in which No-Go trials were added such that there were 16 Go and 16 No-Go trials on each lever (64 total trials/session). In No-Go trials, the lever was presented simultaneously with 2 cues (the house lights turning off, and a small LED light above the lever turning on). A lever press during the 5 second trial caused the lever to retract, the house lights to turn on, the LED light to turn off and a new ITI to begin without any reward for that trial (“Unsuccessful No-Go Trial”). A lack of presses during the 5 sec trial resulted in a reward presentation (“Successful No-Go Trial”). Hit rate was calculated as the proportion of Successful Go trials and the false alarm rate as the proportion of Unsuccessful No-Go trials, respectively averaged over all days. Impulsivity index was calculated for small and large reward levers by subtracting the proportion of correct No-Go trials from the proportion of correct Go trials. This composite index has a maximum score of +1, which indicates highest impulsive responding (always responding on Go and No-Go trials). The minimum score of −1 indicates lowest impulsive behavior, essentially never responding on either No-Go or Go trials. The latency to lever press was also recorded for each trial and averaged across days; the latency was recorded as the maximum 5 seconds if there was no lever press during the trial.

### Statistical Analysis

Group effects were evaluated using analysis of variance (ANOVA), with post hoc Fisher’s PLSD in StatView (SAS Software, Cary, NC) or SPSS (IBM, Armonk, NY). When pairwise comparisons were made following the primary ANOVAs, one-way ANOVAs were first used to test group effects within conditions when there was a significant interaction in the repeated measures ANOVAs, followed by post-hoc Fisher’s LSD if there was an effect of 5-HT_1B_R expression. Two-way repeated measures ANOVAs were used to assess the effects of 5-HT_1B_R (control, no expression, rescued expression) on concurrent choice and devaluation (5-HT_1B_R expression × condition [evaporated milk or standard chow]), random ratio (5-HT_1B_R expression × RR schedule), progressive ratio (5-HT_1B_R expression × 3 days), and extinction (5-HT_1B_R expression × 3 days). For all remaining experiments, 5-HT_1B_R expression levels only included whole life knockout and control. Two-way repeated measures ANOVAs were also used for the standard Go/No-Go (5-HT_1B_R expression × 10 days), delay discounting (5-HT_1B_R expression × delay), and effort discounting (5-HT_1B_R expression × FR schedule). For delay discounting, a linear equation was also fit to data from each mouse. The slope, intercept, and fit (r^2^) were compared between groups using unpaired t-tests. The indifference point, defined as the time delay when the preference was 50% was calculated for each mouse based on the linear equation, and compared between groups using an unpaired t-test. For the lickometer tests of hedonic value/reward reactivity, three-way mixed ANOVAs were used to determine the effects of 5-HT_1B_R expression, sucrose concentration, and fed state (restricted or sated). A two-way mixed ANOVA was used to analyze the effect of CS and genotype on nose poking during the Pavlovian training in the PIT paradigm, and an unpaired t-test was used to compare the difference score between genotypes on the instrumental transfer test. Two-way mixed ANOVAs were used for the effect of reward value on impulsivity in the Variable Value Go/No-Go paradigm validation study (reward size × 10 days for impulsivity index; reward size × trial type for hit rate/false alarm rate and latencies). Three-way mixed ANOVAs were used to assess the effect of reward value and 5-HT_1B_R expression manipulation on impulsivity in the experimental Variable Value Go/No-Go paradigm (5-HT_1B_R expression × reward size × 10 days for impulsivity index; 5-HT_1B_R expression × reward size × trial type for hit rate/false alarm rate and latencies). Mixed ANOVAs, as appropriate, were also used to assess the interaction of sex with these variables. For food consumption, sex was found to have a significant effect, therefore data was analyzed with a three-way mixed ANOVA for the effects of 5-HT_1B_R expression, time, and sex in the restricted condition, and a two-way ANOVA for the effects of 5-HT_1B_R expression and sex in the sated condition.

There were no significant effects of sex on the remaining behaviors measured, and therefore the sexes are combined for all other analyses presented. Results of statistical tests are reported in the figure legends.

## Results

A lack of 5-HT_1B_R expression increases impulsive action, but not impulsive choice. Specifically, mice lacking the 5-HT_1B_ receptor showed increased impulsive action in the Go/No-Go task (Fig 1A), as measured by a reduced ability to inhibit behavioral responding on No-Go trials. While they showed some improvement in their ability to inhibit level presses on No-Go trials over 10 training sessions, this was slower and reduced compared to control mice. We also used a delay discounting paradigm as a second test of impulsivity aimed at measuring the impulsive choice dimension. Interestingly, mice lacking 5-HT_1B_R expression did not show increased impulsivity, but rather an overall *increase* in preference for the large reward (Fig 2B). This is represented by an increased indifference point – the delay length at which small immediate and large delayed rewards are chosen equally, 11.1s (±3.9s) in mice lacking 5-HT_1B_R, compared to 2.5s (±1.0s) in controls. However, the increased preference for the large reward was seen across all delays with no interaction of group and delay, and no group differences in the slope of the discounting function, suggesting that the effect of 5-HT_1B_ is not on impulsivity. Rather, the overall increase in preference for the large reward across all delays is shown by an upward shift in the discounting curve (change in intercept) suggesting that the effect of 5-HT_1B_R on choice may be due to changes in valuation of the reward.

**Figure 1.**
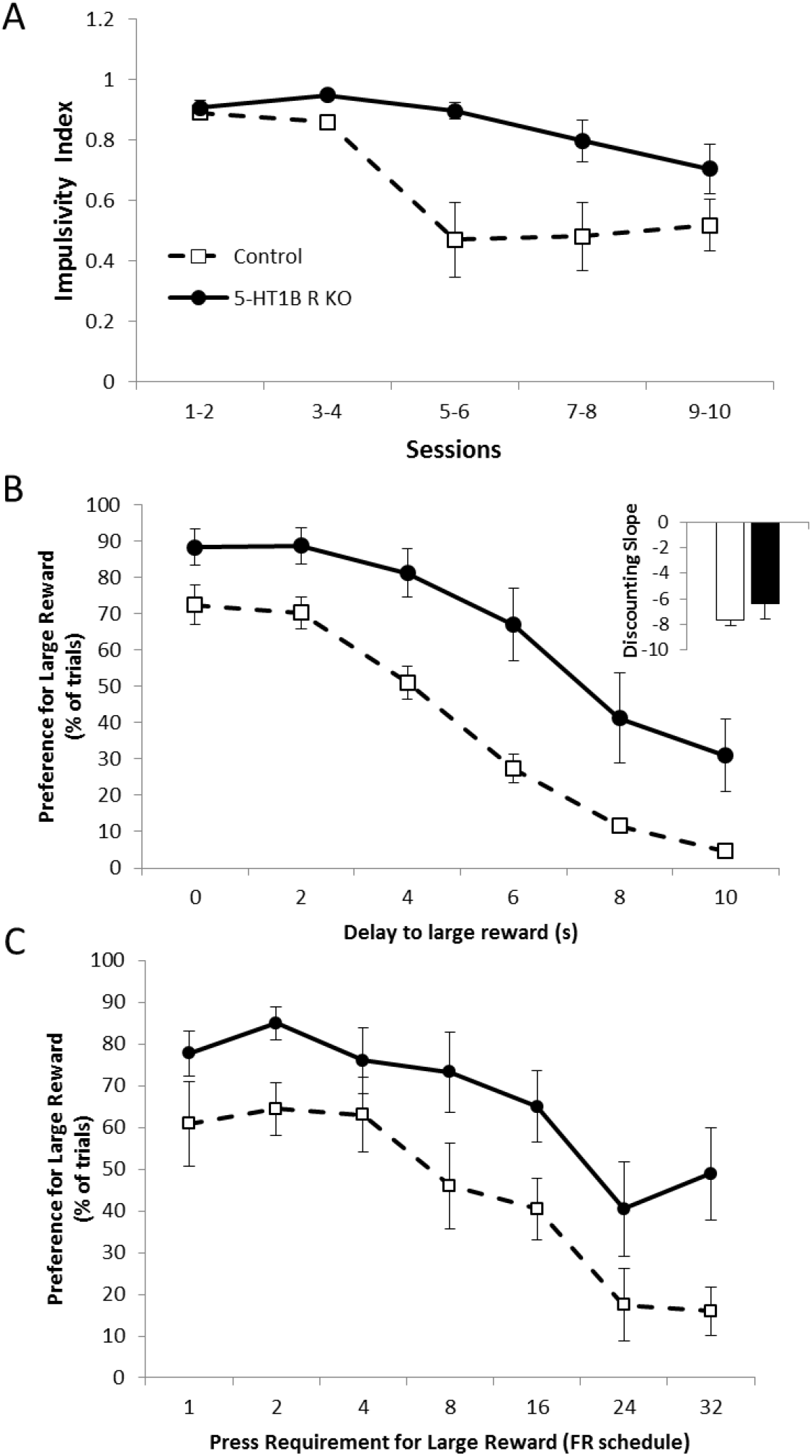
Absence of 5-HT_1B_R increases impulsive action but not delay or effort-based discounting. (A) Impulsivity index calculated as the proportion of successful Go trials minus the proportion of successful No-Go trials is shown as a measure of impulsive action (1.0 is the highest impulsivity that a mouse can display) over 10 days presented in 2-day bins. There were main effects of 5-HT_1B_R expression (F_1,18_=7.0 p<0.05) and session (F_9,162_=8.1, p<0.0001), as well as a 5-HT_1B_R expression × session interaction (F_9,162_=3.1, p<0.005). (B) Data from a delayed discounting paradigm are shown as the percentage of trials on which the large (delayed) reward was chosen, represented over delays ranging from 0 to 10 seconds. Inset shows discounting slope, with more negative slopes indicating a more impulsive choice behavior. There was a significant main effect of delay (F_5,85_=62.6, p<0.001) as well as an effect of 5-HT_1B_R expression on preference for the large reward (F_1,17_=12.4, p<0.005), with no interaction between 5-HT_1B_R expression and delay (F_5,85_=1.6, p>0.05). Additionally, there was no significant difference between groups in indifference point (t_17_=8.6, p<0.05) or discounting slope (t_17_=1.1, p>0.05), though an overall increase in preference for the large reward was seen across all delays in mice lacking 5-HT_1B_R (t_17_=2.0, p=0.058 for intercept). (C) Performance on an effort-based discounting task is shown for mice lacking 5-HT1BR and controls as a percentage of trials in which the large reward was chosen, represented over effort requirements ranging from fixed-ratio (FR)1 to FR32 schedules. There were significant main effects of 5-HT_1B_R expression (F_1, 17_=5.8, p<0.05) and effort requirement (F_6,102_=19.9, p<0.0001) on preference for the large reward, with no significant genotype by schedule interaction (F_6, 102_=0.6, p>0.05). All data are shown as group means +/− SEM.

**Figure 2.**
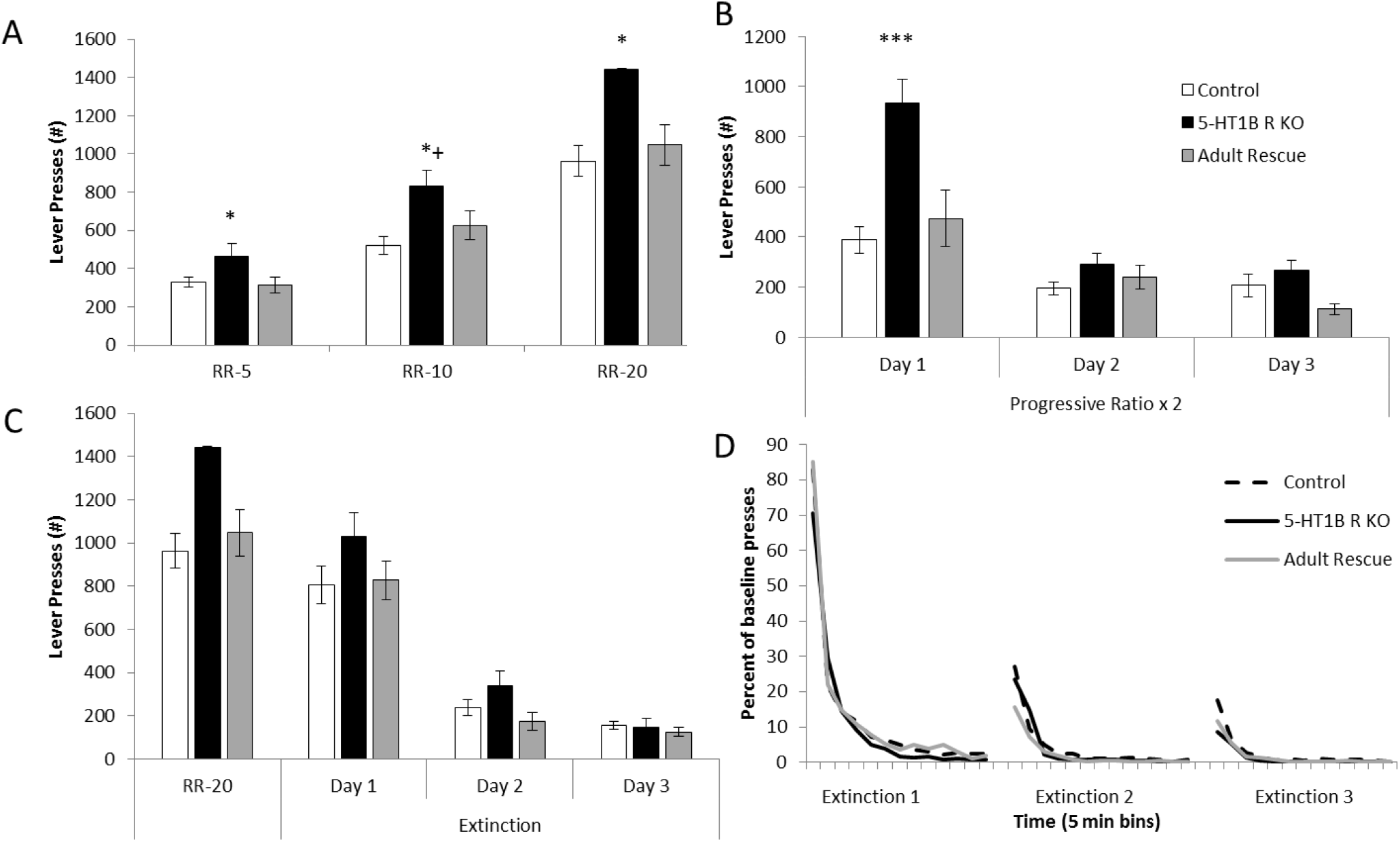
Lack of 5-HT_1B_R increases motivation. (A) Number of lever presses are shown during random ratio 5, 10, and 20 schedules of reinforcement, with main effects of 5-HT_1B_R expression (F_2,55_=8.0, p<0.001) and schedule (F_2,110_=182.4, p<0.0001), and a significant interaction (F_4,110_=3.4, p<0.05). *, p<0.05, 5-HT_1B_R KO compared to Control and Rescue groups for RR5 and RR20. *+, p<0.05, 5-HT_1B_R KO compared to Control group, p=0.054 for 5-HT_1B_R KO vs adult rescue. (B) Number of lever presses are shown for a progressive ratio × 2 schedule of reinforcement, presented over three consecutive days. There were significant main effects of 5-HT_1B_R expression (F_2,53_=7.3, p<0.005) and day (F_2,106_=70.1, p<0.0001) on number of lever presses, and a significant 5-HT_1B_R expression × day interaction (F_4,106_=10.2, p<0.0001). There was a significiant main effect of 5-HT_1B_R expression within Day 1 (F_2,53_=12.0, p<0.0001; ps<0.001 for control and adult rescue vs. 5-HT_1B_R KO, p>0.05 for control vs. adult rescue), but not on Day 2 or 3 (F_2,53_<2.5, p>0.05). ***, p<0.001, 5-HT_1B_R KO compared to Control and Rescue groups. (C) Lever presses shown during 3 extinction sessions, compared to the previous RR-20 session. There was a main effect of time on lever pressing (F_2,110_=153.9, p<0.0001), but no main effect of 5-HT_1B_R expression (F_2,55_=1.5, p>0.05), or interaction of time × 5-HT_1B_R expression (F_4,110_=1.3, p>0.05). (D) Percentage of presses from RR-20 baseline, during 3 sessions of extinction trials, binned by 5 minutes. There was no significant main effect of 5-HT_1B_R expression (F_2,55_=1.6, p>0.05). All data shown are group means +/− SEM.

To assess if the increased preference for the large reward was unique to delays or could be seen more generally in reward value-decision making, we tested the behavior of mice lacking 5-HT_1B_R in an effort-based discounting task. A similar pattern to the delay discounting data emerged – namely that mice lacking 5-HT_1B_R expression showed increased preference for the large reward, over all effort requirements (Fig 1C). As seen in the delay discounting paradigm, there was no significant difference between groups in the slope of the discounting function suggesting that the 5-HT_1B_R doesn’t influence effort-based discounting, but rather might alter baseline reward value scaling.

Given the increased responding for high-effort rewards, we tested the impact of 5-HT_1B_R on motivation. Mice lacking 5-HT_1B_R expression showed increased responding on operant lever pressing tasks including random ratio and progressive ratio schedules, which was also interestingly reversed by adult rescue of receptor expression. For random ratio schedules, 5-HT_1B_R influenced the number of presses - mice lacking 5-HT_1B_R expression pressed 40-50% more than control and adult rescue mice (Fig 2A). As the effort requirements increased, the effect of 5-HT_1B_R expression on lever pressing became larger, again with adult rescue of receptor expression returning behavior to the intact phenotype. The interaction of 5-HT_1B_R expression and schedule suggests that the increased responding is related to goal-directed or motivated responding, rather than a general increase in activity which would likely be read out as increased responding equivalently across all schedules. To assess motivation, we used a progressive ratio schedule of responding. Similar to the random ratio schedules, the absence of 5-HT_1B_R increased lever pressing in PRx2schedule (Fig 2B). Mice lacking 5-HT_1B_R pressed more than controls, which was reversed by adult rescue. Curiously, the effect of 5-HT_1B_ R on lever pressing was only present on Day 1, and not on Days 2 or 3.

One interpretation of the increased responding on the first day, but not subsequent days of testing in the PR is that 5-HT_1B_R KO mice show faster extinction resulting in lower responding after Day 1. To test this idea, we measured extinction of lever pressing behavior in non-rewarded sessions following RR-20 training. Over three days of extinction sessions, while all mice decreased lever pressing, there were no significant effects of 5-HT_1B_R expression on number of lever presses (Fig 2C). Number of lever presses was also normalized to baseline lever pressing behavior to control for the higher starting point in mice lacking 5-HT_1B_R expression, and there were still no differences in extinction rates between groups (Fig 2D). This suggests that the behavioral pattern seen in the progressive ratio task is not due to differences in extinction rate.

To further investigate effort-based decision making, we used a concurrent choice task in which mice were provided with a choice between freely available food/reward in the operant chamber or lever pressing for evaporated milk (Fig 3A). There was an effect of 5-HT_1B_R expression on this effort-based operant task, with mice lacking the receptor continuing to press more despite having a freely available option. While all mice decreased their lever pressing behavior when the freely available option was evaporated milk compared to chow, mice lacking 5-HT_1B_R expression continued pressing at 55% of their baseline rate despite concurrent access to freely available evaporated milk in the operant chamber, while control mice and mice with adult rescue of receptor expression reduced their pressing to 17% and 25% of their baseline rates, respectively. All mice continued to lever press at high rates for evaporated milk when the freely available option was chow (average 97% of baseline). There were no group differences in the consumption of the freely available reward, though all mice consumed more milk than food (Fig 3B). Taken together, these results suggest that 5-HT_1B_R expression could influence either the representation of the outcome value that guides goal-directed action or habitual-like responding.

**Figure 3.**
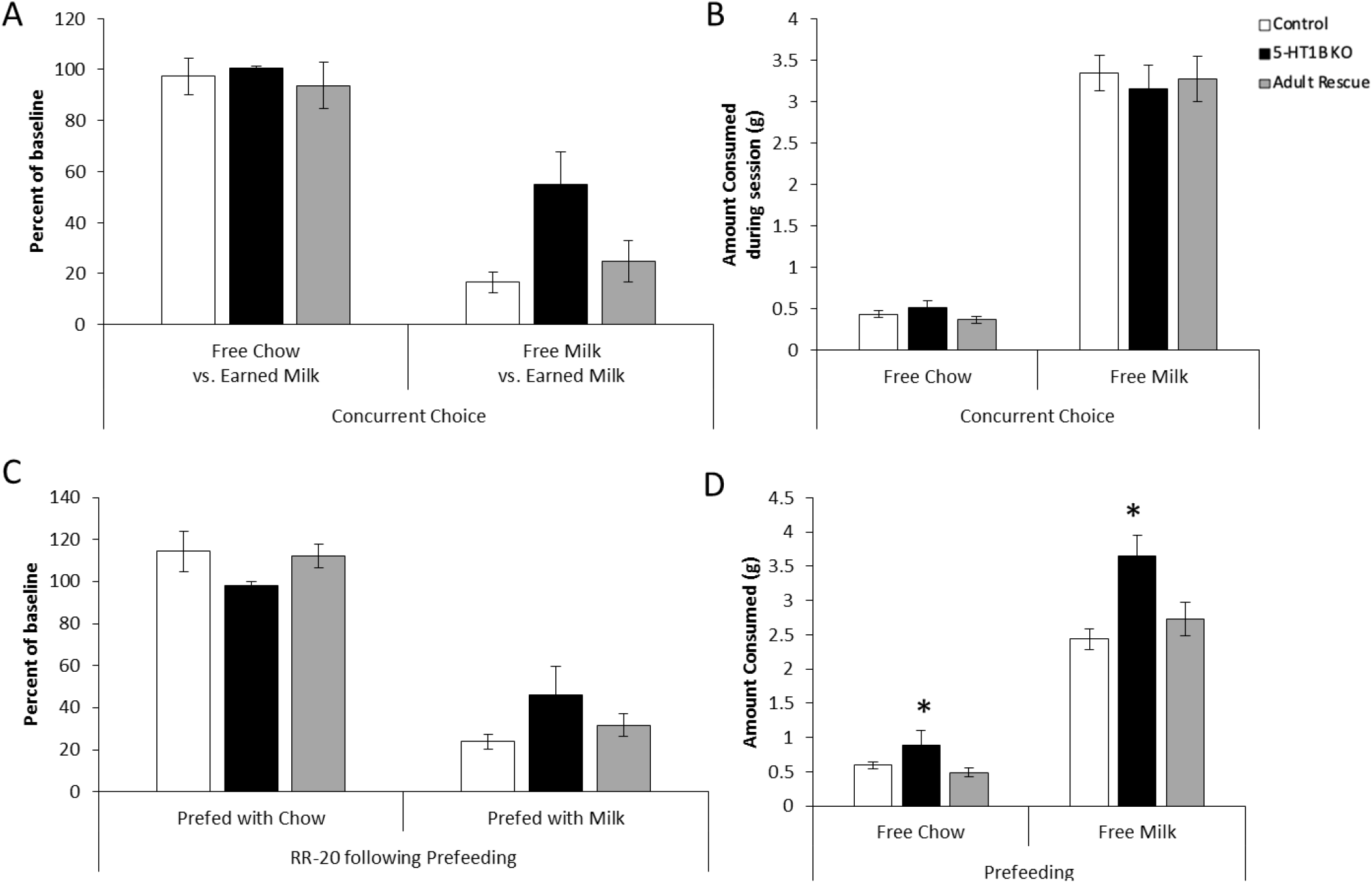
Effects of 5-HT_1B_R on habitual and goal-directed responding. (A) Lever presses are shown as a percentage of a total presses from a baseline RR-20 schedule in conditions in which chow or evaporated milk were presented as free alternatives to lever pressing for evaporated milk. There were main effects of 5-HT_1B_R expression (F_2,55_=3.6, p<0.05) and freely-available option (F_1,55_=106.1, p<0.0001) on lever press behavior, with an interaction term approaching significance (F_2,55_=2.8, p=0.069). (B) The amount of free alternative chow or evaporated milk that was consumed during the operant session is shown, with no group differences (F_2,55_=0.1, p>0.05 for main effect; F_2,55_=0.3, p>0.05 for interaction), though all mice consumed more milk than chow (F_1,55_=325.8, p<0.0001). (C) Lever presses are shown as a percentage of a total presses from a baseline RR-20 schedule in conditions in which mice were prefed chow or evaporated milk before the operant test session, with main effects of prefed option (F_1,38_=89.3, p<0.0001), but no effects of 5-HT_1B_R expression (F_2,38_=0.1, p>0.05 for main effect; F_2,38_=2.1, p>0.05 for interaction). (D) The amount of chow or evaporated milk that was consumed during the prefeeding session prior to operant session is shown. All mice consumed more evaporated milk than chow (F_1,38_=343.9, p<0.0001), but mice lacking 5-HT_1B_R consumed more overall, with a larger effect size in the milk condition (F_2,38_=7.5, p<0.005 for main effect, F_2,38_=5.1, p<0.05 for interaction). *, p<0.05, 5-HT_1B_R KO compared to Control and Rescue groups. All data shown are group means +/− SEM.

We first tested the hypothesis that mice lacking 5-HT_1B_R respond more habitually and are less guided by the outcome/goal of their actions. To do this, we measured goal-directed behavior following satiety-induced devaluation of the reward. There were no significant effects of 5-HT_1B_R expression in this test of habitual-like behavior. All mice similarly reduced responding when pre-fed with evaporated milk reward, but not when pre-fed with chow, showing that mice were responding in a similar goal-directed, rather than habitual manner on the RR-20 schedule. Furthermore, this suggests that the increased lever pressing behavior in mice lacking 5-HT_1B_R in the concurrent choice paradigm is likely not a function of increased habitual-like responding, but rather potentially due to altered representations of the reward value. This is also supported by an increased intake in the pre-operant test satiety induction with mice lacking 5-HT_1B_R expression consuming more reward in the pre-operant feeding sessions, with the increase being larger in the milk compared to the chow condition (Fig 3D). Overall these results suggest that the behavioral differences seen in mice lacking 5-HT_1B_R are not likely due to increased habitual-like responding.

Next, we addressed the hypothesis that an exaggerated representation of outcome value could arise from a difference in hedonic reactions to the reward. First we measured amount of consumption to varied concentrations of evaporated milk reward used in operant tests (Fig S1). We found that mice lacking 5-HT_1B_R consume more evaporated milk than controls, and increase their consumption more as the reward concentration goes up suggesting that they scale reward value differently. Next, in order to test the effect of 5-HT_1B_R on hedonic value more directly, we used a standard lickometer to examine licking behavior to different concentrations of sucrose [49,50]. The lickometer reduces some of the motivational components required for operant-based tasks, and also eliminates the contribution of post-ingestive factors found in consumption tests, therefore allowing measurement of a more immediate hedonic reaction through analysis of licking behavior. Mice lacking 5-HT_1B_R expression showed increased in hedonic reactivity, as measured by increased licking for sucrose compared to controls (Fig 4A,B). Across different conditions, mice lacking 5-HT_1B_R also showed greater increases in total licks as the motivational state or value increased suggesting that these mice were scaling reward value differently. We also examined lick rate during the first two minutes of the sessions to remove any potential confound of effects of satiety on hedonic readouts since brief access durations reduce non-taste effects [46,47]. Mice lacking 5-HT_1B_ R expression showed increased lick rates that approached significance (Fig 4C, D). Together, these results show that mice lacking 5-HT_1B_R expression have exaggerated licking responses compared to controls across all conditions, though importantly maintain the normal relative changes based on motivational state and concentration. This suggests that 5-HT_1B_R acts to shift the scale of the normal valuation of reward.

**Figure 4.**
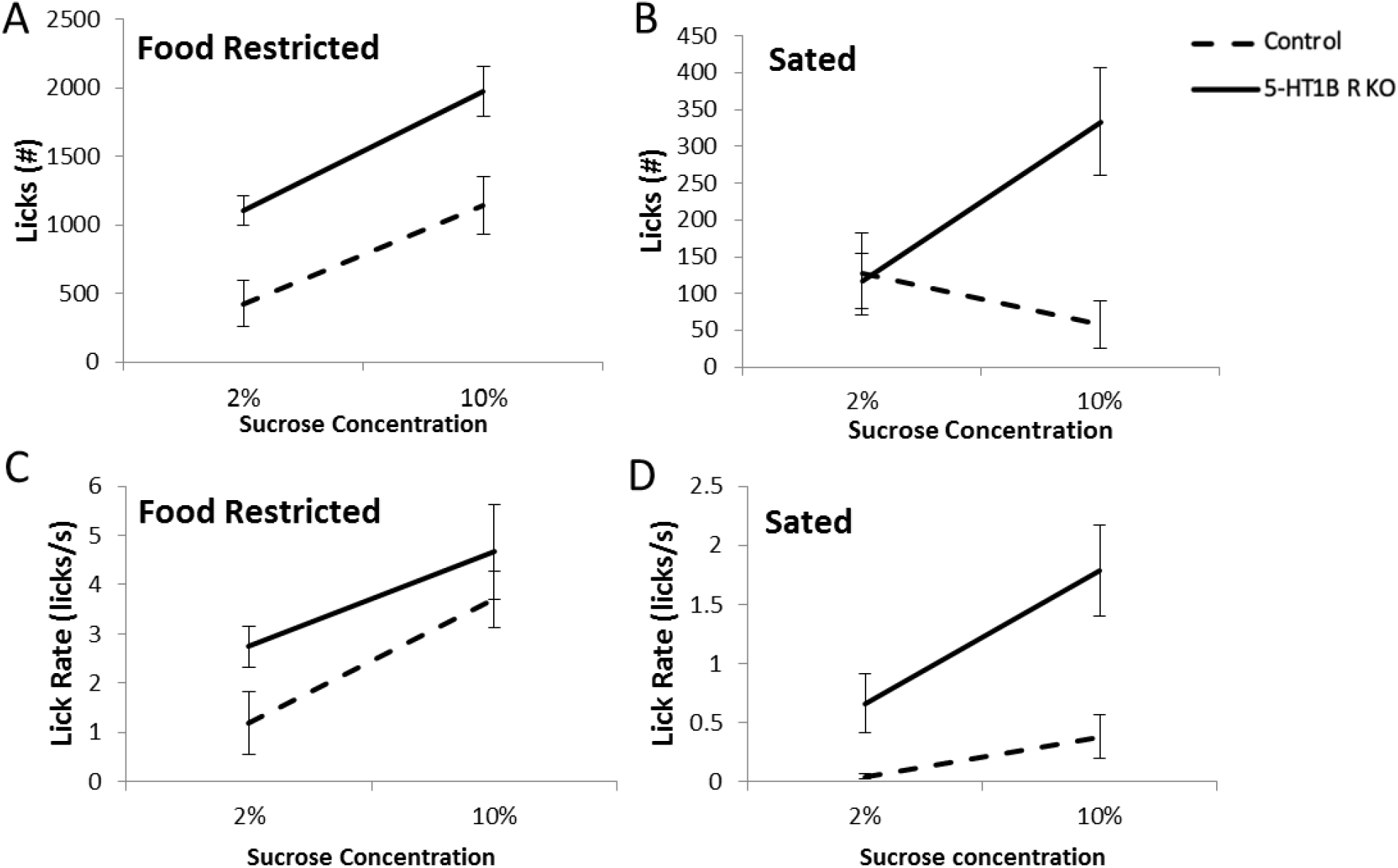
5-HT_1B_R expression influences hedonic valuation. (A, B) Total number of licks to a spout delivering sucrose, shown in food restricted (A) and sated (B) conditions to 2% and 10% sucrose, is greater in mice lacking 5-HT_1B_R (F_1,9_=12.0, p<0.01). There were also main effects of deprivation state (food restricted vs. sated; F_1,9_=89.9, p<0.001) and sucrose concentration (2% vs. 10%, F_1,9_=76.7, p<0.001), as well as a fed state × concentration (F_1,9_=85.9, p<0.001), with mice licking more in the restricted and 10% sucrose conditions. There was also a significant interaction of 5-HT_1B_R expression × fed state (F_1,9_=8.6, p<0.05) and a suggestive interaction of 5-HT_1B_R expression × concentration (F_1,9_=4.8, p=0.056). (C, D) Lick rate in the first 2 minutes of the session, for food restricted (C) and sated (D) conditions to 2% and 10% sucrose. There was a main effect of 5-HT_1B_R expression approaching significance (F_1,9_=4.3, p=0.069), with 5-HT_1B_R KOs showing higher lick rates overall compared to controls. All groups had increased lick rates toward 10% sucrose (F_1,9_=35.8, p<0.001 for main effect) and in the restricted condition (F_1,9_=41.2, p<0.001 for main effect), with the increase being the largest in the restricted 10% condition (F_1,9_=11.5, p<0.01 for concentration × fed state interaction). There were no interactions with 5-HT_1B_R expression for lick rates (all ps>0.05). All data are shown as group means +/− SEM.

We also addressed the possibility that the effects of 5-HT_1B_R expression were due to increased feeding drive rather than specific to reward responsivity. There were no significant differences in chow intake between 5-HT_1B_R KO and littermate control mice in either restricted (Fig S2A) or sated conditions (Fig S2B). These results suggest that the increase in reward-motivated behaviors seen in the absence of 5-HT_1B_Rs is not due to a general increase in hunger or feeding, and thus lends support to our interpretation that 5-HT_1B_R influences the valuation of palatable rewards.

So far, we’ve shown that mice lacking 5-HT_1B_R respond more vigorously to palatable rewards. We suggest that the exaggerated hedonic responses may be the result of higher value representations of reward, which also serve to increase goal-directed behavior relative to controls. We performed a Pavlovian-to-instrumental transfer (PIT) study to assess if a reward-paired cue motivates instrumental responding differently in the absence of 5-HT_1B_R expression. During the initial associative learning phase, all mice learned to discriminate between cues as measured by increased head entries into reward receptacle during the CS+ compared to the CS− (Fig 5A). There were no significant effects of genotype during the Pavlovian training phase. However, mice lacking 5-HT_1B_R expression displayed higher levels of lever pressing for the CS+ in the instrumental transfer test compared to controls (Fig 5B). Importantly the transfer test was performed in the absence of any prior instrumental training, highlighting the role of the CS+ in promoting acquisition of instrumental behavior, rather than a potentiation of a previously learned response-outcome association. This suggests that the value attributed to the CS+ motivates instrumental responding more in the absence of 5-HT_1B_R expression.

**Figure 5.**
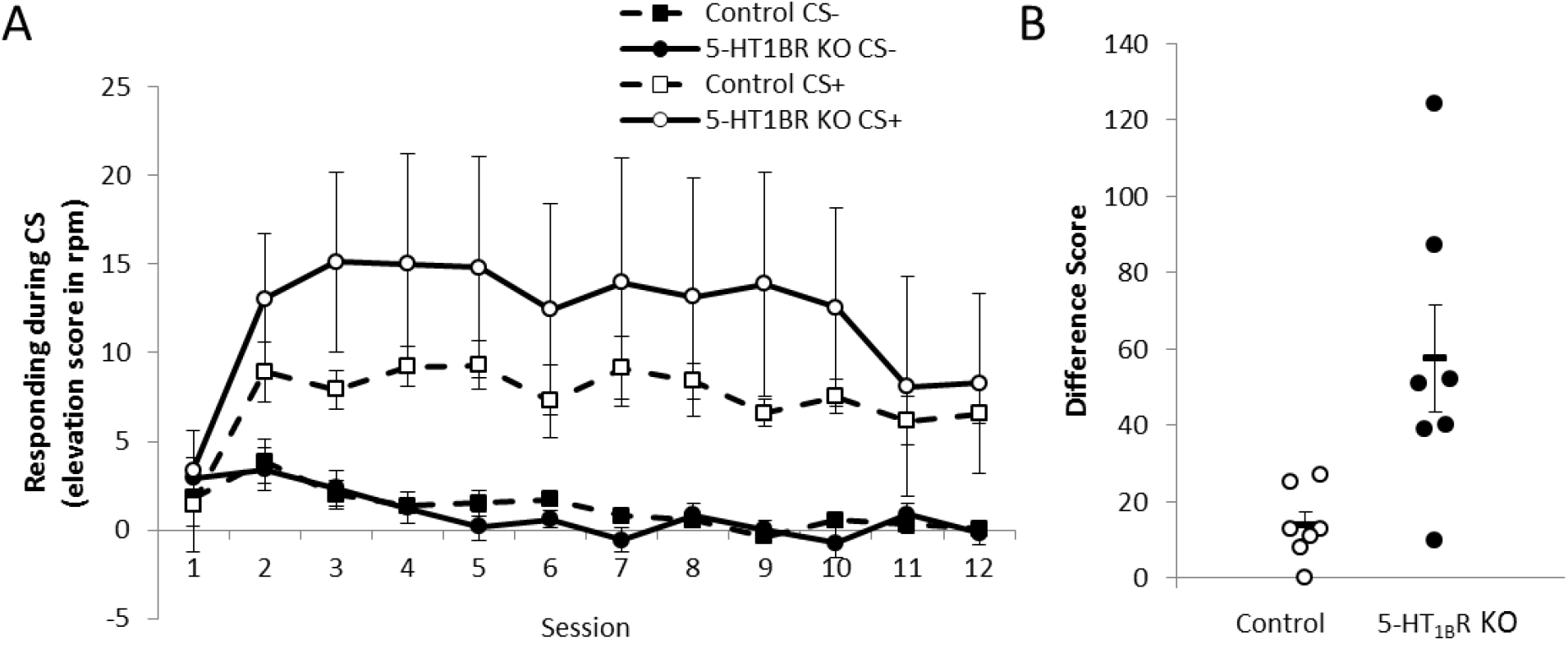
Decreasing reward value ameliorates 5-HT_1B_R-related impulsivity. (A) Schematic shows the Variable Value Go/No-Go paradigm used to assess the effect of reward value on impulsive action. (B) Impulsivity index calculated as the proportion of successful Go trials minus the proportion of successful No-Go trials is shown as a measure of impulsive action (1.0 is the highest impulsivity that a mouse can display) over 10 days presented in 2-day bins. Data is shown for each small and large reward trials for Controls and 5-HT_1B_R KOs, with main effects of reward size (F_1,16_=25.9, p<0.001) and 5-HT_1B_R expression (F_1,16_=6.3, p<0.05). (C) Hit rate for Go trials and (D) false alarm rate for No-Go trials again had significant main effects of reward size (F_1,16_=25.9, p<0.001) and 5-HT_1B_R expression (F_1,16_=6.3, p<0.05). (E) Latency to press the lever for Go trials and for (F) No-Go trials was also influenced by reward size (F_1,16_=46.7, p<0.001) and 5-HT_1B_R expression (F_1,16_=4.7, p<0.05). All data are shown as group means +/− SEM.

Given the effects of 5-HT_1B_ on driving reward associated cue-motivated behavior, we next tested if alterations in reward valuation could also explain the increased impulsivity seen in mice lacking 5-HT_1B_R expression. We first developed and tested a novel paradigm – the Variable Value Go/No-Go, to directly examine how manipulating reward value could impact impulsive action on a trial-by-trial basis (Fig 6A). By varying reward value in a Go/No-Go paradigm within a single session, we could compare impulsivity within mice between trials in which large or small rewards were expected. In control mice, we found that mice were more impulsive on large compared to small reward trials (Fig S3). Using this novel paradigm, we were then able to investigate the role of increased reward valuation in 5-HT_1B_R-induced deficits in impulsive action, and test whether the increased impulsivity in mice lacking 5-HT_1B_R expression could be ameliorated by decreasing reward value. Our results show that the increased impulsivity in mice lacking 5-HT_1B_R was ameliorated, in part, by reducing the reward value by three times. Specifically, behavior on small reward trials in mice lacking 5-HT_1B_R was similar to that of high reward trials in controls (Fig 6B). Overall, both a lack of 5-HT_1B_R expression and a larger reward magnitude increased impulsivity as seen in false alarm rates and hit rates (Fig 6C,D). These effects could also be read out by decreased response latencies in mice lacking 5-HT_1B_R and in controls on large reward trials (Fig 6E,F). Interestingly this shows that 5-HT_1B_R-associated impulsivity can be reduced by decreasing the reward value and suggests that alterations in reward value alone can lead to increased impulsivity. Our data suggest that reward reactivity is an important behavioral component to measure in the study of the neural circuits underlying impulsivity, and point to a behavioral mechanism through which serotonin influences impulsive action.

**Figure 6.**
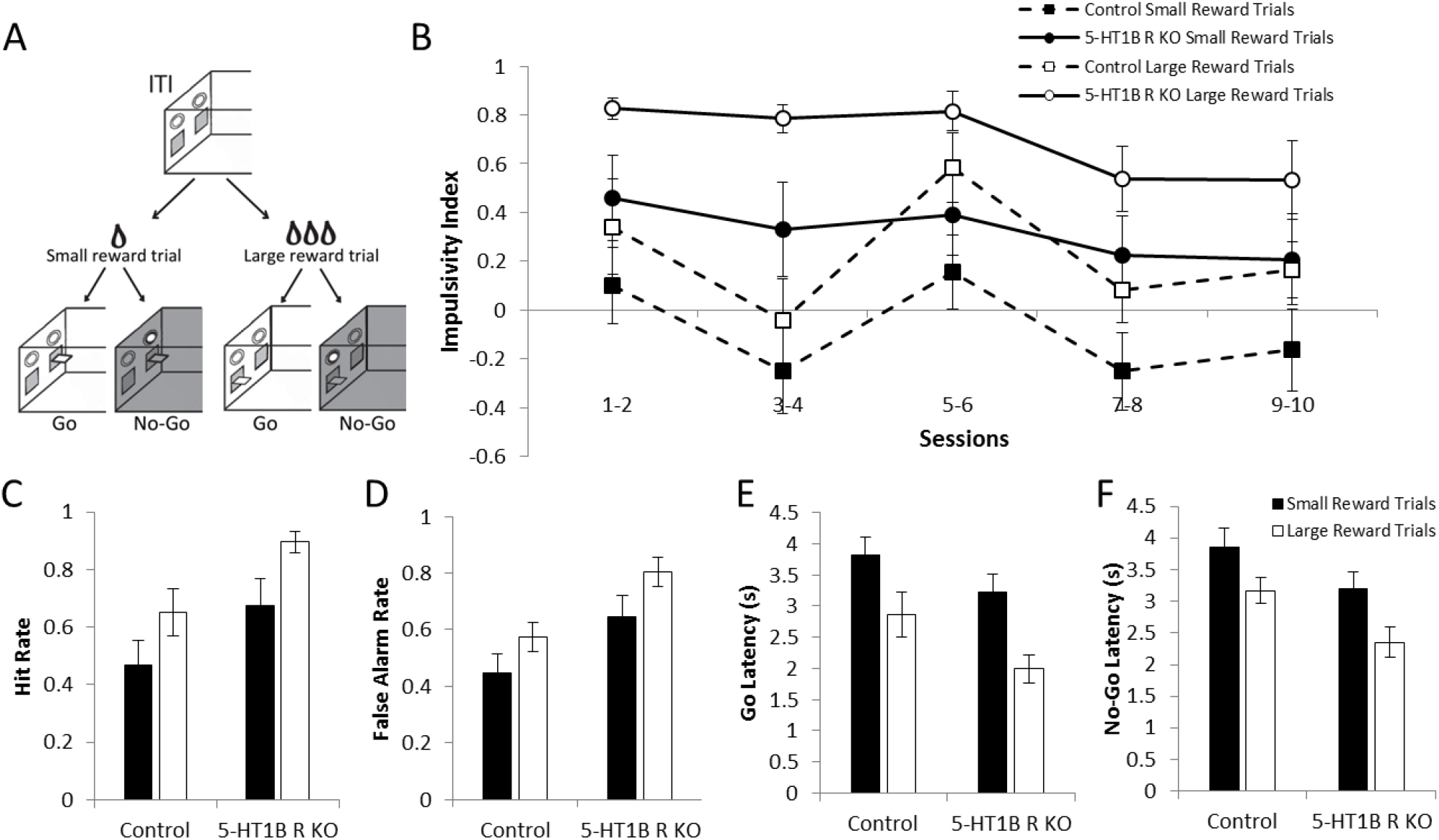
Decreasing reward value ameliorates 5-HT_1B_R-related impulsivity. (A) Diagram of Variable Value Go/No-Go paradigm. (B) Impulsivity index calculated as the proportion of successful Go trials minus the proportion of successful No-Go trials is shown as a measure of impulsive action (1.0 is the highest impulsivity that a mouse can display) over 10 days presented in 2-day bins. Data is shown for each small and large reward trials for controls and 5-HT_1B_R KOs, with main effects of reward size (F_1,16_=25.9, p<0.001) and 5-HT_1B_R expression (F_1,16_=6.3, p=0.023). (C) Hit rate for Go trials and (D) false alarm rate for No-Go trials again had significant main effects of reward size (F_1,16_=25.9, p<0.001) and 5-HT_1B_R expression (F_1,16_=6.3, p=0.023). (E) Latency to press the leverfor Gotrials and for (F) No-Gotrials was also influenced by reward size (F_1,16_=46.7, p<0.00l) and 5-HT_1B_R expression (F_1,16_=4.7, p=0.046). All data are shown as group means +/− SEM.

## Discussion

Overall, our data points to a role for altered reward value representation in the serotonin modulation of impulsive behavior. Specifically, we show that 5-HT_1B_R expression influences goal-directed behavior, motivation, PIT, and hedonic valuation, along with effects on impulsive action, but not impulsive choice. We also tested a subset of these phenotypes with adult rescue of 5-HT_1B_R expression which rescued normal behavior, suggesting ongoing modulation of neural circuits rather than compensatory effects. While, we have previously shown that 5-HT_1B_R receptor expression influences impulsive action during adulthood, we now provide a behavioral mechanism of action. First, mice lacking 5-HT_1B_R expression show increases in hedonic responses to sucrose, compared to controls. We propose that this may be a readout of increased valuation of rewards. This interpretation is consistent with a recent study which illustrates the influence of 5-HT_1B_R on the representation of outcomes through changes in sensitivity to the sensory qualities of reinforcers [51]. Interestingly, in our studies mice lacking 5-HT_1B_R expression also show increased goal-directed responding, which is sensitive to extinction and devaluation, and we propose that this is driven by increased valuation of the reward. This is supported by increased responding seen in the PIT study in 5-HT_1B_R KO mice, which suggests that a higher attribution of value to the CS+ (acquired during the Pavlovian training) motivates higher levels of instrumental responding. Taken together with the effects of 5-HT_1B_R on impulsive action, these data point to the possibility that the influence of 5-HT_1B_R on reward valuation may contribute to the effects on goal-directed behavior and motivation, as well as on impulsive action.

Previous studies in humans and animal models have examined the relationship between hedonic value and impulsivity [14,52,53]. In rats, increased sucrose-seeking is associated with increased impulsive action (measured in the 5-choice serial reaction time task) [54], and rats bred for high sucrose consumption displayed higher levels of impulsive action (on the Go/No-Go task) when responding for cocaine [53], and also higher levels of impulsive choice (on the delay discounting task) [55]. Though in humans, one study showed that increased hedonic value measured with varying sweet concentrations is associated with increases in impulsive choice (assessed in a delay discounting task), but not impulsive action (measured in a Go/No-Go paradigm) [14]. However, a confound in the interpretation of many of these studies suggesting associations between reward value and impulsivity arises from between-subjects designs measuring more trait-like phenotypes. This leaves open the possibility for another trait-level behavioral construct to mediate the association between reward value and impulsivity (e.g. learning about appetitive goal-directed behavioral contingencies). In order to test the causal association of higher valued incentive stimuli leading to increased impulsivity, we developed a within-subject, within-session experiment varying reward value, and could therefore directly measure the effects on impulsive action in the Go/No-Go task. The results from this Variable Value Go/No-Go paradigm show that increased reward value causes increased impulsive action as measured by a decrease in behavioral inhibition in No-Go trials. This supports a causal role for reward value in impulsive action. Furthermore, we were able to increase the impulsivity in controls to similar levels to that seen in mice lacking 5-HT_1B_R by tripling the reward value. This suggests that the impulsive phenotype seen in mice lacking 5-HT_1B_R could feasibly be derived by only changing the subjective value of the reward.

Given that past studies have implicated serotonin in the regulation of feeding and locomotion, alternative interpretations for our data include that the phenotypes are driven by an influence of 5-HT_1B_R on increased hunger drive or general activity. To rule out hunger, we directly measured feeding behavior, and found no effect on food intake in fed or restricted conditions. Importantly, past work implicating 5-HT_1B_R in body weight regulation in the original 5-HT_1B_R knockout mouse line (generated in the 1980s) includes a methodological limitation of not controlling for genetic background (using non-congenic, non-littermate controls) making reported effects on bodyweight difficult to interpret [56,57]. Other pharmacology work has reported that 5-HT_1B_R agonists decrease food consumption, however we suggest that these effects are derived from a non-specific behavioral effect on motivation, and because of the use of large doses that may bind non-specifically [57]. For example, the authors report no effect on feeding at 5mg/kg of the 5-HT_1B_R agonist CP-94,253, a dose that elicits behavioral effects on impulsivity, and report that the effects on food intake were only seen at doses more than twice at high (10-20mg/kg). These higher doses also have suggestive or significant effects on feeding in 5-HT_1B_R KO mice suggesting non-specific binding. Additionally, the idea that 5-HT_1B_R influences motivation for non-food reward is further supported by past studies showing increased motivation for cocaine in 5-HT_1B_R KO mice [25]. To address the possibility that the reported phenotypes are due to hyperactivity, we also referenced past work in the 5-HT_1B_R KO mouse. This past report of 5-HT_1B_R involvement in modulating a hyperactive response was specific to non-entrained stimuli (unexpected intruder in the resident-intruder task or disturbance by an experimenter) in a startle-like manner rather than a conditioned response to an entrained stimulus [58]. We would argue the unexpected and potentially stressful stimuli which induce the startle-like hyperactivity are unlike any stimuli presented in our studies. In fact, our data shows increased responding in a stimulus-free well-learned action-outcome contingency. Additionally, our results show that extinction and pre-feeding both reduce responding to control levels which would not be expected in a model of general hyperactivity. Based on these reports and our results, we maintain our initial interpretation that the behavioral effects seen here are not likely due to a change in feeding drive or activity, but rather an exaggeration of the representation of hedonic value of rewarding stimuli.

Past work has examined the role of 5-HT_1B_R in the modulation of a number of models of psychiatric disorders which present with reward-related dysfunctions including addiction and depression. However, there has been limited careful exploration of the underlying behavioral mechanisms that contribute to these 5-HT_1B_R-associated phenotypes which is important for our understanding of complex behavioral processes found across multiple psychiatric disorders. Our data presents a basic reward reactivity-related phenotype that may serve as a framework for synthesizing these previously reported varied effects. Additionally, our work on behavioral mechanisms adds value to past and future studies investigating the neural circuit mechanisms through which 5-HT_1B_R exerts its effects on more complex phenotypes seen in substance use disorder and major depressive disorder.

Early studies in the original 5-HT_1B_R KO mouse line showed an increased motivation for [25,59,60] and decreased motivation for cocaine following 5-HT_1B_ receptor agonist administration [26], now we can investigate the circuit specific effects of 5-HT_1B_R on drug taking behavior. We propose a role for 5-HT_1B_Rs expressed on the terminals of nucleus accumbens shell neurons, particularly in the rewarding properties of low-doses of cocaine that don’t induce reward behavior in controls [61–63]. Additional work also implicates 5-HT_1B_R expression on these accumbens projection neurons in the consumption of ethanol [64]. Alterations in ventral striatal 5-HT_1B_R expression are also seen in major depressive disorder [65,66], and in rodents, 5-HT_1B_R expression in the ventral striatum is implicated in inducing depressive states [67,68]. Finally, the 5-HT_1B_R has also been examined in the context of social reward. Particularly, the rewarding properties of social behavior in mice requires activation of 5-HT_1B_Rs in the nucleus accumbens [69]. On one hand, these neural mechanisms point to a potential neural circuit mechanism for our results, and concurrently, our studies provide the behavioral mechanistic link between the circuit level mechanisms and the complex behavioral readouts.

It is interesting to note that the increasing reward value was associated with increased impulsive action, but not impulsive choice. Specifically, in the delay discounting task used to measure impulsive choice, mice lacking 5-HT_1B_R expression chose the large delayed reward more than controls across all delays. This increased preference was seen in trials without any delay, suggesting that the differences seen in the delay discounting task are not due to changes in the tolerance to delay, but rather to some factor that is common across all delays, such as reward valuation. This interpretation is consistent with past studies which have found that rats prone to attribute incentive salience to reward cues show increased impulsive action but not impulsive choice [13]. Though there is evidence that also supports a link between the sensitivity to the hedonic valuation of sweet reward and impulsive choice, it is possible that 5-HT_1B_R signaling acts through striatal mechanisms to link reward value and impulsive action rather than cortical areas like the vmPFC which may mediate the link between sweet taste activated reward and delay discounting [14,70,71]. This would fit with the lower relative levels of 5-HT_1B_R protein expression in the cortex compared to the ventral striatum [72,73]. Indeed, a lack of 5-HT_1B_R expression results in increased dopamine release in the nucleus accumbens, which is a substrate for goal-motivated behaviors and impulsive action [74–77]. Understanding the 5-HT_1B_R-induced changes in reward reactivity that correlate with behavioral inhibition in impulsive action paradigms, but not temporal discounting in impulsive choice paradigms, may shed light on the neural circuits which underlie psychopathologies that have disordered reward responsiveness and impulsivity.

Overall, we propose that a behavioral mechanism for the effect of serotonin signaling on impulsive action is alterations in reward reactivity. While prior work demonstrated a role for 5-HT_1B_R expression in the modulation of impulsivity as well as the rewarding properties of drugs and social stimuli, our studies provide a unifying hypothesis for all of these effects by identifying a common underlying behavioral substrate. Specifically, we show that there is a causal effect of reward value on impulsive action in our novel Variable Value Go/No-Go paradigm and that decreasing reward value alone is enough to decrease 5-HT_1B_R-associated impulsivity. These studies contribute to research aimed at understanding factors that contribute to increases in impulsivity seen in clinical populations. Additionally, our research points to the utility of serotonin receptor-specific treatment strategies to alter hedonic valuation for psychiatric disorders which involve dysregulated impulsivity.

---

**Supplemental Figure 1.**
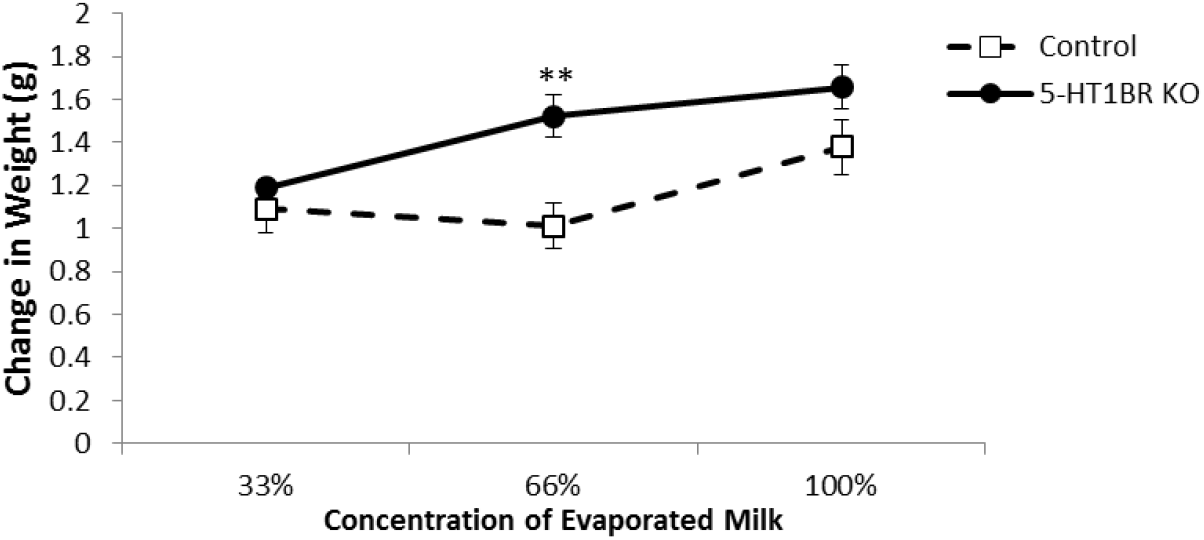
Lack of 5-HT_1B_R does not alter food consumption. (A) Food (standard chow) consumed in restricted (over 1, 3, and 24 hours) and (B) sated (over 24 hours) conditions, in Control and 5-HT_1B_R KO mice. There were no significant group differences in food intake in either restricted (F_1,14_=0.4, p>0.05 for 5-HT_1B_R expression; F_2,28=_0.3, p>0.05 for 5-HT_1B_R expression × time interaction) or sated conditions (F_1,14_=0.4, p>0.05). Mice consume more food over longer periods of time in a deprived state (F_2,28_=463.8, p<0.001), with males consuming more in general and increasing with length of time (F_1,14_=10.3, p<0.01 for sex; F_2,28_=14.6, p<0.001 for sex × time interaction). This effect of increased consumption in males also occurred in the 24h sated period (F_1,14_=11.9, p<0.005). Importantly, there was no interaction of 5-HT_1B_R expression and sex in either experiment (all ps>0.05). All data are shown as group means +/− SEM.

**Supplemental Figure 2.**
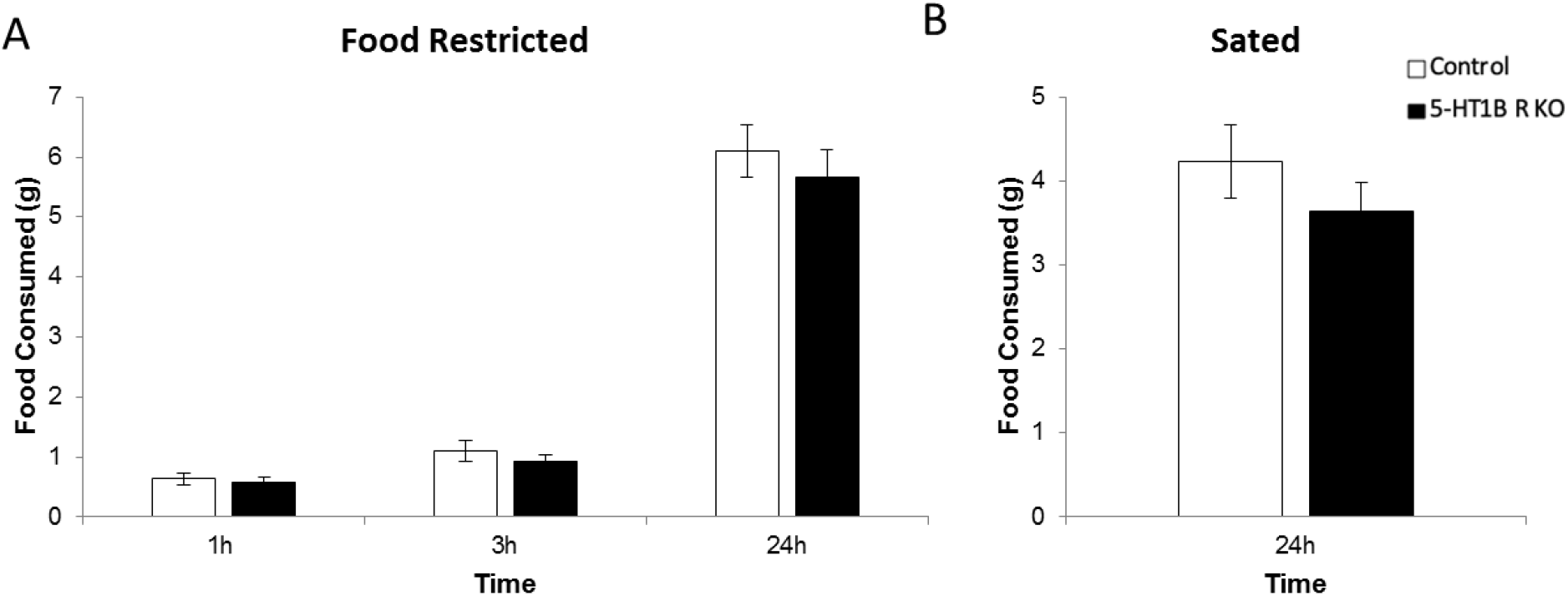
Lack of 5-HT_1B_R increases reward consumption. Change in bodyweight following 5 minute free access to 33%, 66%, and 100% evaporated milk in Control and 5-HT_1B_R KO mice. All mice showed increased consumption of evaporated milk as the concentration increased (F_2,34_= 15.2, p<0.001), mice lacking 5-HT_1B_R showed increased consumption of evaporated milk compared to controls (F_1,17_=5.9, p<0.05). A significant interaction of concentration and 5-HT_1B_R expression suggests that the mice lacking the 5-HT_1B_R showed a greater reactivity to the quality of the reward (F_2,34_= 4.4, p<0.05). This increase was significant at the intermediate 66% concentration (**, p<0.01). All data are shown as group means +/− SEM.

**Supplemental Figure 3.**
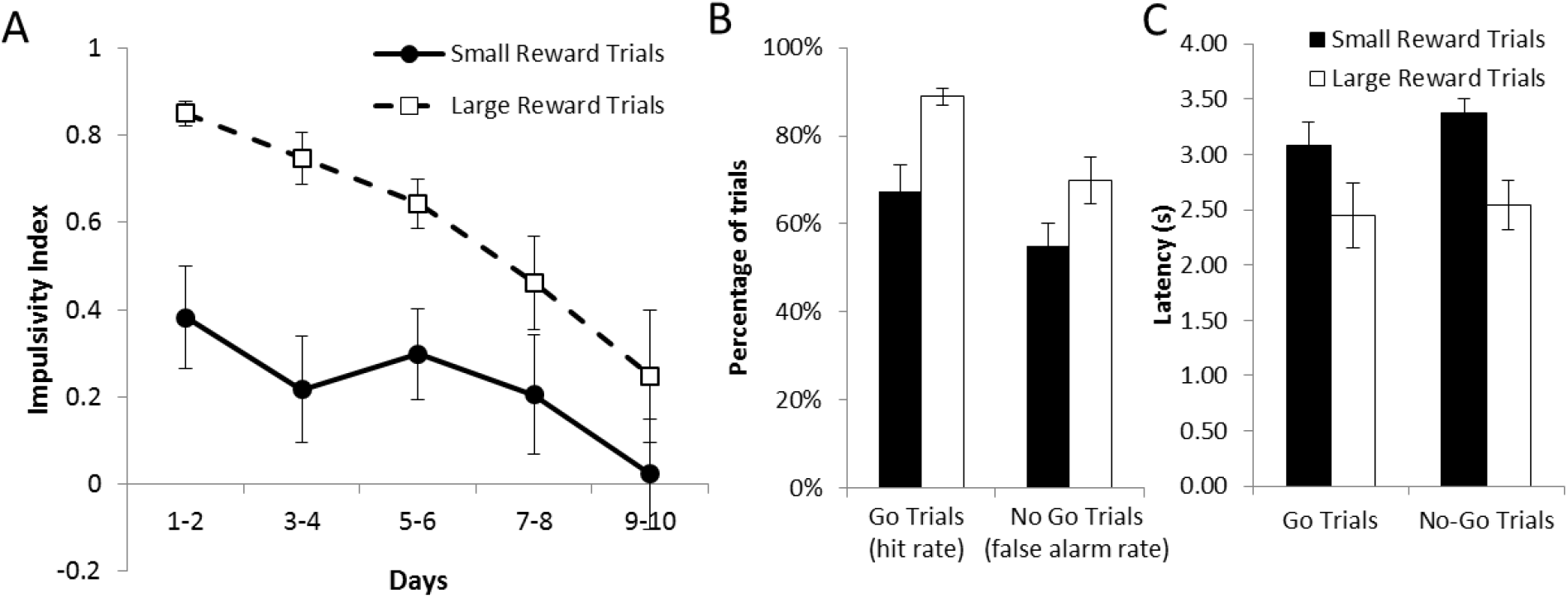
Reward value influences impulsive action on a trial-by-trial basis. (A) Impulsivity index calculated as the proportion of successful Go trials minus the proportion of successful No-Go trials is shown as a measure of impulsive action (1.0 is the highest impulsivity that a mouse can display) over 10 days presented in 2-day bins. Data is shown for each small and large reward trials. Mice are more impulsive for larger rewards (F_1,11_=19.1, p<0.001 for main effect; F_9,99_=2.2, p<0.05 for interaction), while impulsivity in general decreases over sessions (F_9,99_=7.0, p<0.001). (B) Hit rate for Go trials and false alarm rate for No-Go trials. The increased impulsivity index in the large reward condition was influenced by both more correct Go trials and more incorrect No-Go trials (F_1,11_=19.1, p=0.001 for main effect of reward size) (C) Latency to press the lever for Go trials and No-Go trials, with faster responding on large reward trials (F_1,11_=9.5, p=0.01). All data are shown as group means +/− SEM.

**Supplemental Table 1.**
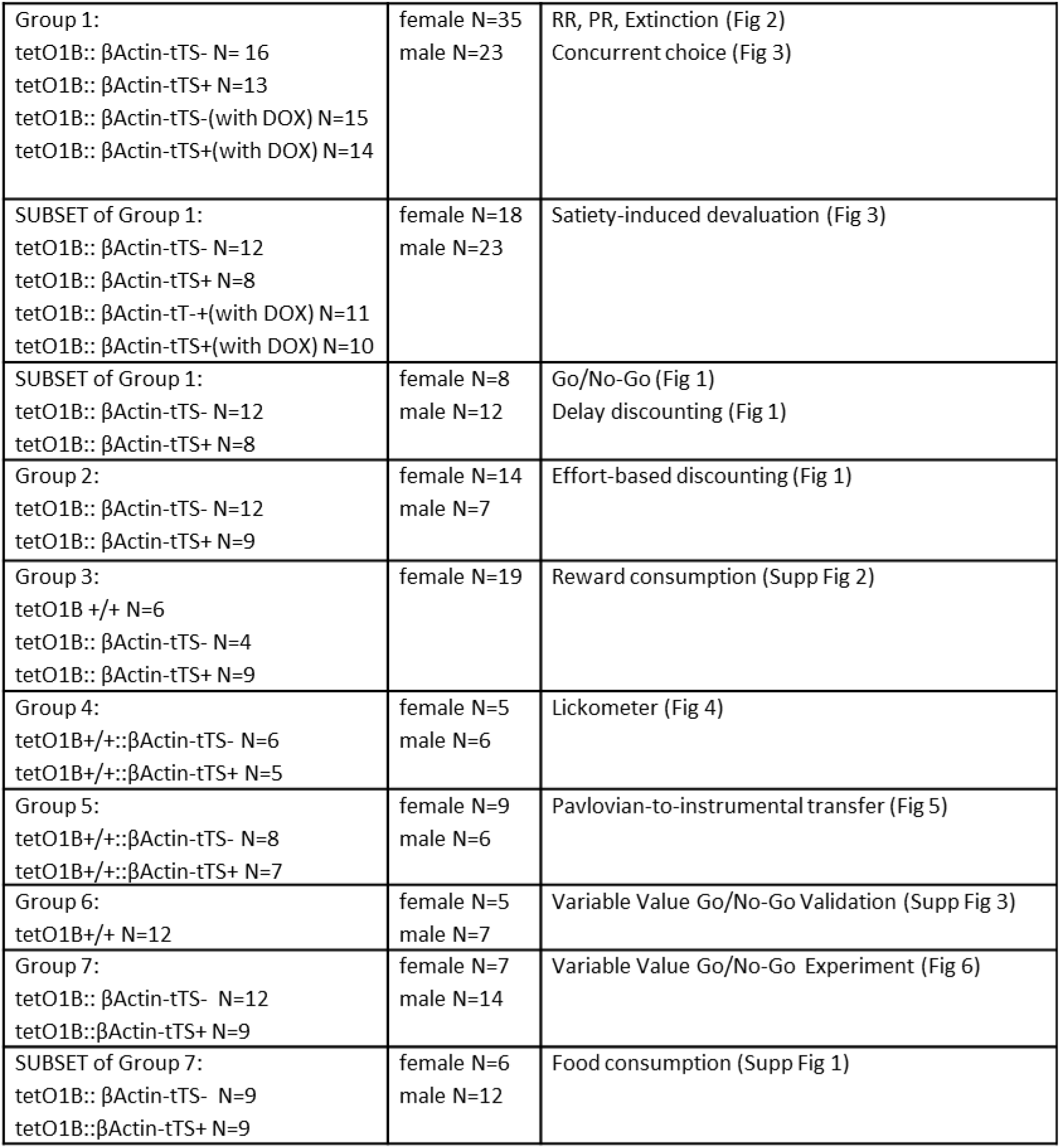

## Acknowledgments

We would like to thank Jun Ho Lee for lab management, Abigail Kalmbach for helpful assistance with the lickometer set-up, and Min Oh, Arati Sharma, and Anne George for help with data collection. We would also like to thank Professor Robert Leaton for his helpful comments on a previous version of this manuscript.

## Author Contributions

SSD, EL, VMM, and KN acquired and managed the data. SSD, EL, and KN conducted data analysis. SSD and KN wrote the first draft of the paper. SSD, KN, and PB designed the studies. All authors contributed to and approved the final version of the paper.

